# Mitochondrial CircRNA CircMT-RNR2 Safeguards Antioxidant Defense to Support Fibroblast Functions in Wound Repair

**DOI:** 10.1101/2025.09.03.673954

**Authors:** Guanglin Niu, Jennifer Geara, Yongjian Chen, Yanwei Xiao, Zhuang Liu, Pehr Sommar, Aoxue Wang, Xiaowei Zheng, Ning Xu Landén

## Abstract

Diabetic foot ulcers (DFUs) are a debilitating diabetes complication in which mitochondrial dysfunction and oxidative stress are prominent but mechanistically unresolved features. Here, we identify the mitochondria-encoded circular RNA circMT-RNR2 as a novel modulator of mitochondrial redox homeostasis in human skin wound healing. CircMT-RNR2 is reduced in DFU patient tissue and diabetic mouse wounds, enriched in dermal fibroblasts, and localized to mitochondria. Its loss impairs fibroblast proliferation, migration, extracellular matrix production, and contraction by destabilizing the mitochondrial antioxidant protein PRDX3, leading to elevated oxidative stress, mitochondrial damage, and mitophagy. In murine and human *ex vivo* wound models, circMT-RNR2 knockdown delays healing, whereas overexpression accelerates repair and boosts antioxidant defenses. These findings position circMT-RNR2 as a mitochondrial guardian of skin healing and a promising therapeutic target for DFUs.

**One Sentence Summary:** CircMT-RNR2, a mitochondria-encoded circular RNA suppressed in diabetic foot ulcers, promotes fibroblast function and maintains mitochondrial redox balance via stabilization of the antioxidant protein PRDX3, offering a promising therapeutic target for chronic wound repair.

## INTRODUCTION

Diabetes mellitus is a widespread chronic metabolic disorder with severe complications, among which diabetic foot ulcers (DFUs) are particularly debilitating (*1*). DFUs are characterized by chronic inflammation, impaired vascularization, and stalled wound healing, often associated with elevated reactive oxygen species (ROS) levels that exacerbate tissue damage (*2*). Despite medical advances, effective DFU therapies remain limited, highlighting the need to better understand their underlying mechanisms (*3*).

Mitochondria are central to cellular metabolism, ATP production, and redox signaling (*4, 5*). Mitochondrial redox balance is critical for wound repair, as controlled ROS signaling promotes fibroblast proliferation, extracellular matrix (ECM) synthesis, and tissue remodeling, whereas excessive ROS causes oxidative damage, chronic inflammation, and impaired healing (*6*). In DFUs, this balance is disrupted by mitochondrial dysfunction and weakened antioxidant defenses, including downregulation of peroxiredoxin III (PRDX3), which amplifies oxidative stress and stalls repair (*7–9*). Restoring mitochondrial redox homeostasis therefore represents a promising strategy for improving chronic wound outcomes.

Circular RNAs (circRNAs), which are covalently closed single-stranded RNAs, have emerged as key regulators of gene expression by modulating transcription and splicing, stabilizing mRNAs, influencing translation, and interfering with signaling pathways (*10*). While most circRNAs originate from the nuclear genome, recent studies have identified circRNAs encoded by the mitochondrial genome (mecciRNAs) in animals (*11–13*). Although their biogenesis and turnover remain unclear, mecciRNAs are increasingly recognized as regulators of mitochondrial function (*14–16*). For example, mecciND1 and mecciND5 act as molecular chaperones to facilitate mitochondrial protein import (*11*). MecciRNA SCAR binds ATP5B to inhibit mitochondrial permeability transition pore opening and reduce ROS output, preivously shown to influence conditions such as chronic lymphocytic leukemia and nonalcoholic steatohepatitis (*17, 18*). Given the central role of mitochondrial dysfunction in DFUs, we hypothesized that mecciRNAs may also contribute to diabetic wound pathogenesis.

Here, we identify circMT-RNR2, a mecciRNA downregulated in DFUs, as a critical regulator of fibroblast function and wound repair. CircMT-RNR2 maintains mitochondrial redox homeostasis by interacting with and stabilizing the antioxidant protein PRDX3, thereby limiting ROS-induced damage and promoting fibroblast proliferation, extracellular matrix deposition, and tissue contraction. These findings reveal a novel link between mecciRNAs and redox control in diabetic wound healing, highlighting a promising therapeutic target for chronic wounds.

## RESULTS

### CircMT-RNR2 expression is downregulated in diabetic foot ulcer

From our previous RNA-sequencing analysis of circRNAs in human skin and wound tissues, we identified a set of mecciRNAs (*19*). Among these, circMT-RNR2—a circular RNA encoded by the mitochondrial gene *MT-RNR2*—was significantly reduced in DFU tissue (n = 65) compared with normal human skin (n = 35), as confirmed by qRT-PCR (**Fig. 1A, Table S1**). In both skin and DFU samples, qRT-PCR revealed higher circMT-RNR2 expression in the dermis than in the epidermis (**Fig. 1B**), consistent with its enrichment in dermal fibroblasts relative to epidermal keratinocytes (**Fig. 1C**). Based on these observations, we focused subsequent experiments on human adult dermal fibroblasts (HDFa) to investigate circMT-RNR2 function.

**Figure 1.**
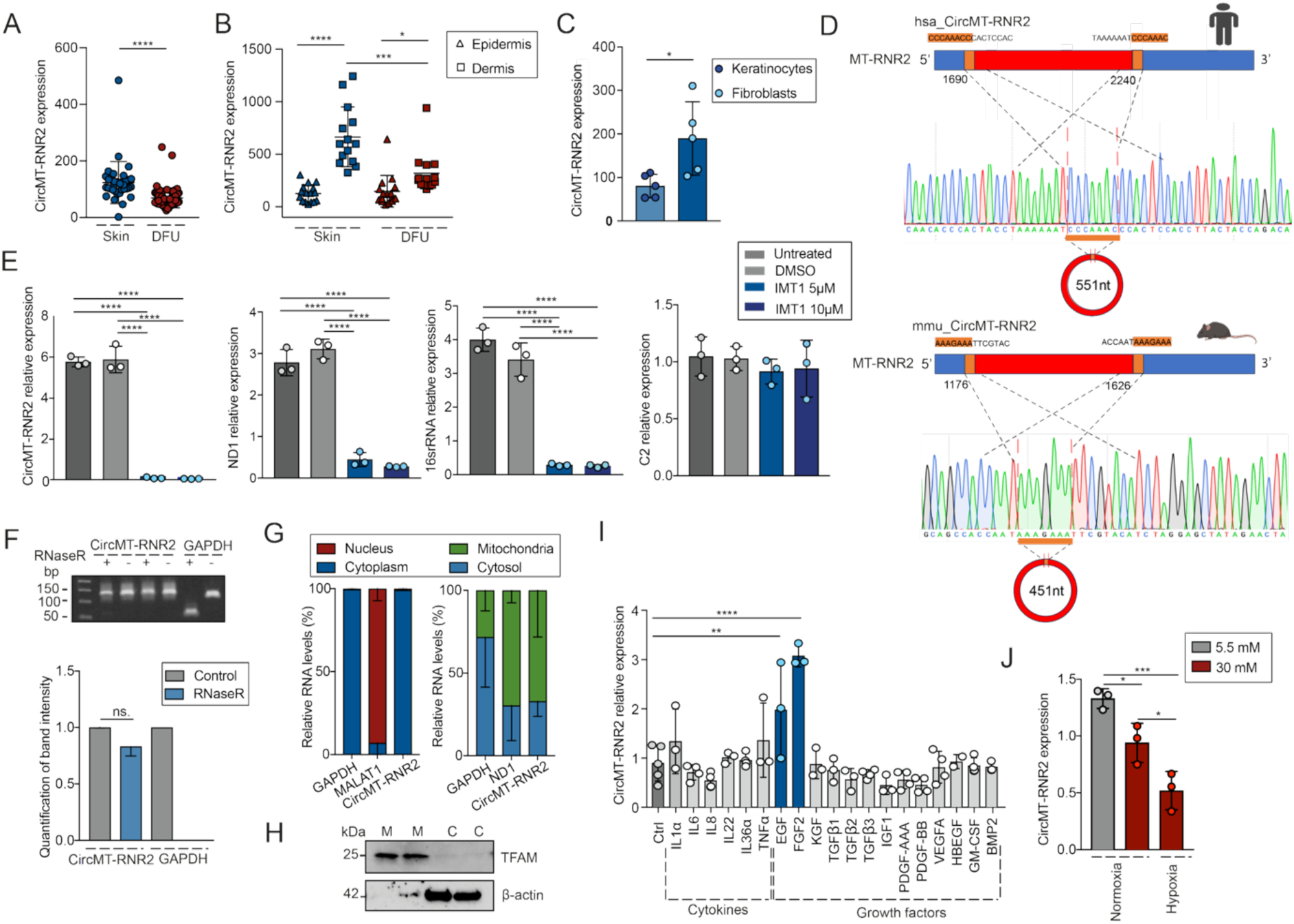
CircMT-RNR2 expression is downregulated in diabetic foot ulcer. qRT-PCR of circMT-RNR2 expression in full-thickness skin biopsies from healthy donors (n=35) and diabetic foot ulcer (DFU) patients (n=65) **(A)**, and in the epidermal and dermal layers of skin (n=16) and DFU samples (n=16) **(B)**. **(C)** qRT-PCR analysis of circMT-RNR2 expression in Keratinocytes and Fibroblasts isolated from human skin (n=5). **(D)** Genomic location and junction sites of circMT-RNR2. Sanger sequencing confirms human (up) and mouse (down) sequences; flanking repeats in orange. **(E)** qRT-PCR of circMT-RNR2, mitochondrial genes (*ND1*, 16S rRNA), and nuclear gene (C2) after POLRMT inhibition (IMT1, 5 or 10 µM, n=3). **(F)** Agarose gel electrophoresis of circMT-RNR2 and *GAPDH* RT-PCR products from RNase R–treated vs. control RNA in HDFa cells; Bar graph shows band quantification. **(G)** qRT-PCR of circMT-RNR2, *GAPDH*, *MALAT1* and *ND1* in nuclear, cytoplasmic, mitochondrial and cytosolic fractions of HDFa cells (n=2). **(H)** Western blot of mitochondrial and cytosolic fractions confirming purity (TFAM: mitochondrial marker; β-actin: cytosolic marker). M = mitochondria, C = cytosol. **(I)** qRT-PCR of circMT-RNR2 in HDFa treated with cytokines or growth factors (24 h, n= 3-5). **(J)** circMT-RNR2 expression in HDFa under low (5.5 mM) or high (30 mM) glucose, with or without hypoxia (n=3). ns: not significant, *P<0.05, **P<0.01, ***P<0.001, ****P<0.0001, paired Student t-test, or unpaired Student t-test, or one-way ANOVA and Turkey’s multiple comparisons test.

The junction site of circMT-RNR2 was validated by Sanger sequencing. As reported for other mecciRNAs (*11, 13*), we identified repetitive sequence motifs flanking the junction— CCCAAAC in human and AAAGAAA in mouse—potentially involved in mecciRNA biogenesis (**Fig. 1D**). The mitochondrial origin of circMT-RNR2 was supported by the observation that treatment of HDFa with IMT1, a mitochondrial RNA polymerase inhibitor, significantly reduced circMT-RNR2 abundance. This decrease paralleled reductions in mitochondrial RNAs (*ND1* and 16S rRNA) but not the nuclear transcript *C2* (**Fig. 1E**). The circular structure of circMT-RNR2 was further confirmed by its resistance to RNase R digestion, which degrades linear but not circular RNAs (**Fig. 1F**) (*20*).

To determine the subcellular localization of circMT-RNR2, we fractionated HDFa into nuclear, cytoplasmic, and mitochondrial compartments. Enrichment of *MALAT1* (nuclear), *GAPDH* (cytoplasmic), and *ND1* (mitochondrial) confirmed fractionation efficiency (**Fig. 1G**), which was further validated at the protein level by Western blot using TFAM (mitochondrial) and β-actin (cytosolic) markers (**Fig. 1H**). These results showed that circMT-RNR2 is predominantly localized to mitochondria in human dermal fibroblasts (**Fig. 1G**).

We next investigated wound microenvironmental factors that might regulate circMT-RNR2 expression. Treatment of HDFa with cytokines and growth factors essential for wound repair revealed that Epidermal Growth Factor (EGF) and Fibroblast Growth Factor 2 (FGF2) significantly increased circMT-RNR2 levels (**Fig. 1I**). In contrast, high glucose and hypoxic conditions reduced its expression (**Fig. 1J**). Together, these findings suggest that the reduced circMT-RNR2 observed in DFU may result from a combination of low EGF/FGF2 levels (*21, 22*), hyperglycemia, and hypoxia characteristic of this chronic wound type.

### Loss of circMT-RNR2 impairs fibroblast functions critical for wound healing

We next examined the role of circMT-RNR2 in fibroblast-driven wound repair processes, including proliferation, migration, contraction, and ECM production (*23*). Since mitochondria harbor RNA interference machinery (*24–26*) and siRNA-mediated knockdown of mitochondrial circRNAs has been demonstrated (*26*), we silenced circMT-RNR2 in HDFa cells using siRNAs targeting its junction site (si-CircMT-RNR2). qRT-PCR confirmed efficient mitochondrial knockdown of circMT-RNR2 without affecting linear *MT-RNR2* expression (**Fig. S1A)**.

Knockdown of circMT-RNR2 impaired several fibroblast functions critical for wound repair. Incucyte live-cell imaging revealed a marked reduction in proliferation, which was further supported by decreased expression of the proliferation marker *MKI67* (**Fig. 2A–B**). In scratch-wound assays, circMT-RNR2–deficient fibroblasts displayed significantly delayed migration compared to controls (**Fig. 2C)**. Furthermore, circMT-RNR2 knockdown blunted TGF-β1– induced expression of ECM genes (*COL1A1, FN1, ELN*) as well as *ACTA2*, which encodes α-SMA, a key driver of fibroblast contractility (**Fig. 2D**).

**Figure 2.**
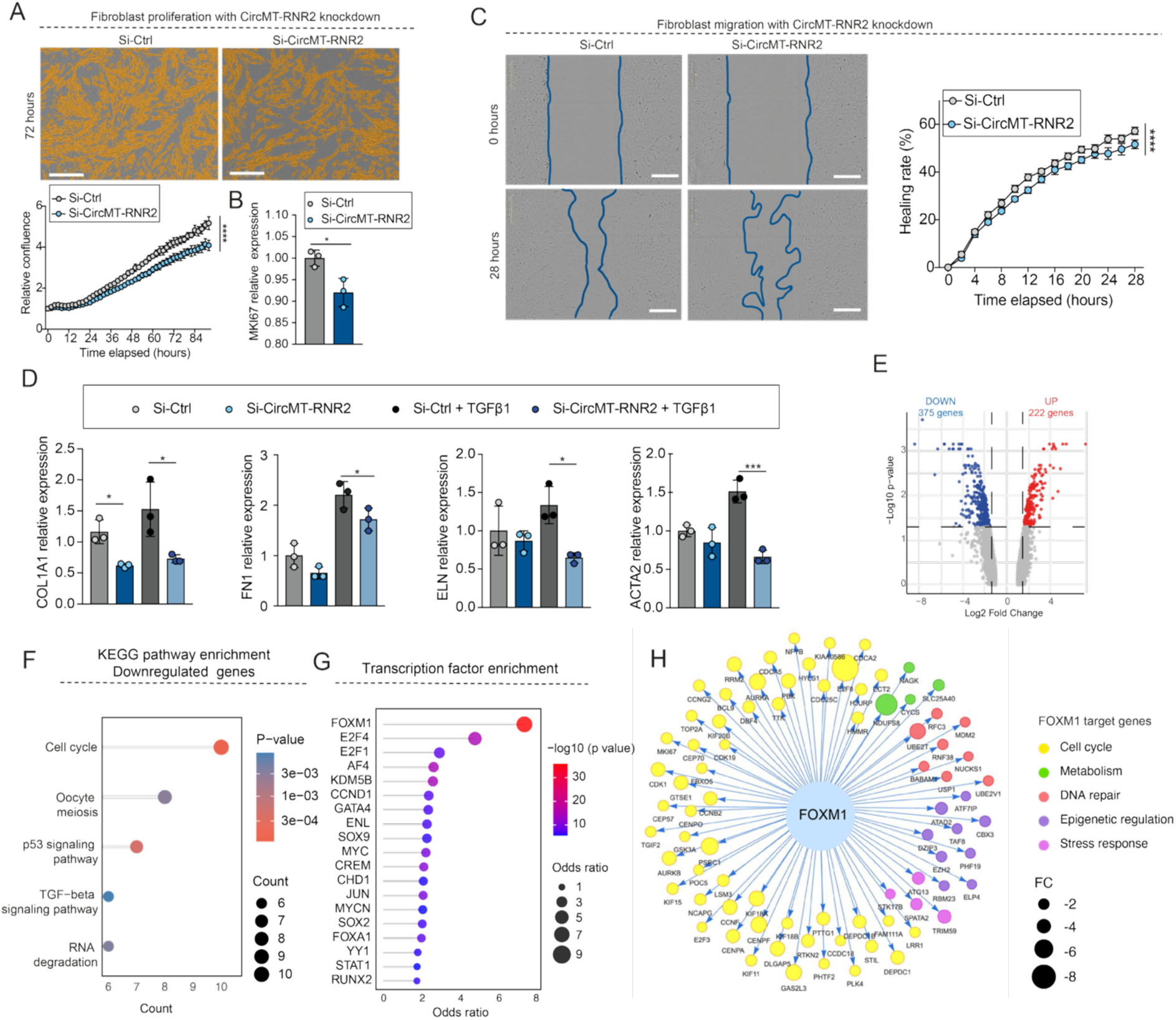
Loss of circMT-RNR2 impairs fibroblast functions critical for wound healing. **(A)** Proliferation assay of HDFa cells treated with si-Ctrl or si-circMT-RNR2 with quantification of relative confluence (n=6) ; scale bar, 400 µm. **(B)** *MKI67* expression in HDFa cells with circMT-RNR2 knockdown (microarray analysis, n=3). **(C)** Scratch-wound migration assay over 28 h in HDFa cells with si-Ctrl or si-circMT-RNR2, with quantification of wound closure rate (%); scale bar, 400 µm. **(D)** qRT-PCR of *COL1A1*, *FN1*, *ELN* and *ACTA2* in HDFa with Si-Ctrl or Si-CircMT-RNR2, with or without TGF-β1 stimulation (24 h, n=3). **(E)** Volcano plot of differentially expressed genes between si-Ctrl and si-CircMT-RNR2 (375 downregulated in blue, 222 upregulated in red; thresholds: FDR < 0.05, |log2FoldChange| > 1.5). KEGG pathway **(F)** and transcription factor (**G**) enrichment of downregulated genes. **(H)** FOXM1 target genes; node color indicates functional categories, node size reflects Fold Change (FC). *P < 0.05, **P < 0.01, ***P < 0.001, ****P < 0.0001 (unpaired Student t-test, or two-way ANOVA and multiple comparisons).

Transcriptomic microarray analysis revealed 375 downregulated and 222 upregulated genes following circMT-RNR2 knockdown (FDR < 0.05, |log2FoldChange| > 1.5) (**Fig. 2E, Table S2**). KEGG pathway enrichment showed that the cell cycle was the most significantly affected pathway among the downregulated genes (**Fig. 2F, Table S3**). Transcription factor (TF) enrichment analysis further identified FOXM1 as the top upstream regulator of these genes, with most FOXM1 targets belonging to the cell cycle pathway (**Fig. 2G–H, Table S4, S5**). Given that FOXM1 is a well-established regulator of fibroblast migration, oxidative stress response, inflammation, and wound repair—and is implicated in DFU pathology (*27–29*)— these findings suggest that circMT-RNR2 supports fibroblast function at least in part by maintaining FOXM1-driven transcriptional programs. Overall, circMT-RNR2 emerges as a critical regulator of fibroblast proliferation, motility, ECM production, and contraction, thereby promoting effective wound healing.

### CircMT-RNR2 safeguards mitochondrial redox homeostasis

Given its mitochondrial origin and localization, we next investigated whether circMT-RNR2 influences mitochondrial function. Microarray analysis following circMT-RNR2 knockdown identified 16 downregulated and 9 upregulated mitochondrial genes based on the Human MitoCarta3.0 database (**Fig. 3A**) (*30*). Notably, downregulated genes included *GSR*, *PRDX2*, *CISD1,BCL2L13, PPIF, SLC25A40,* and *CYCS,* which were all involved in the response to oxidative stress pathway (*31–37*) (**Fig. 3B**). Consistent with this, gene set enrichment analysis (GSEA) revealed significant enrichment of the oxidative stress response pathway among downregulated genes (**Fig. 3C**).

**Figure 3.**
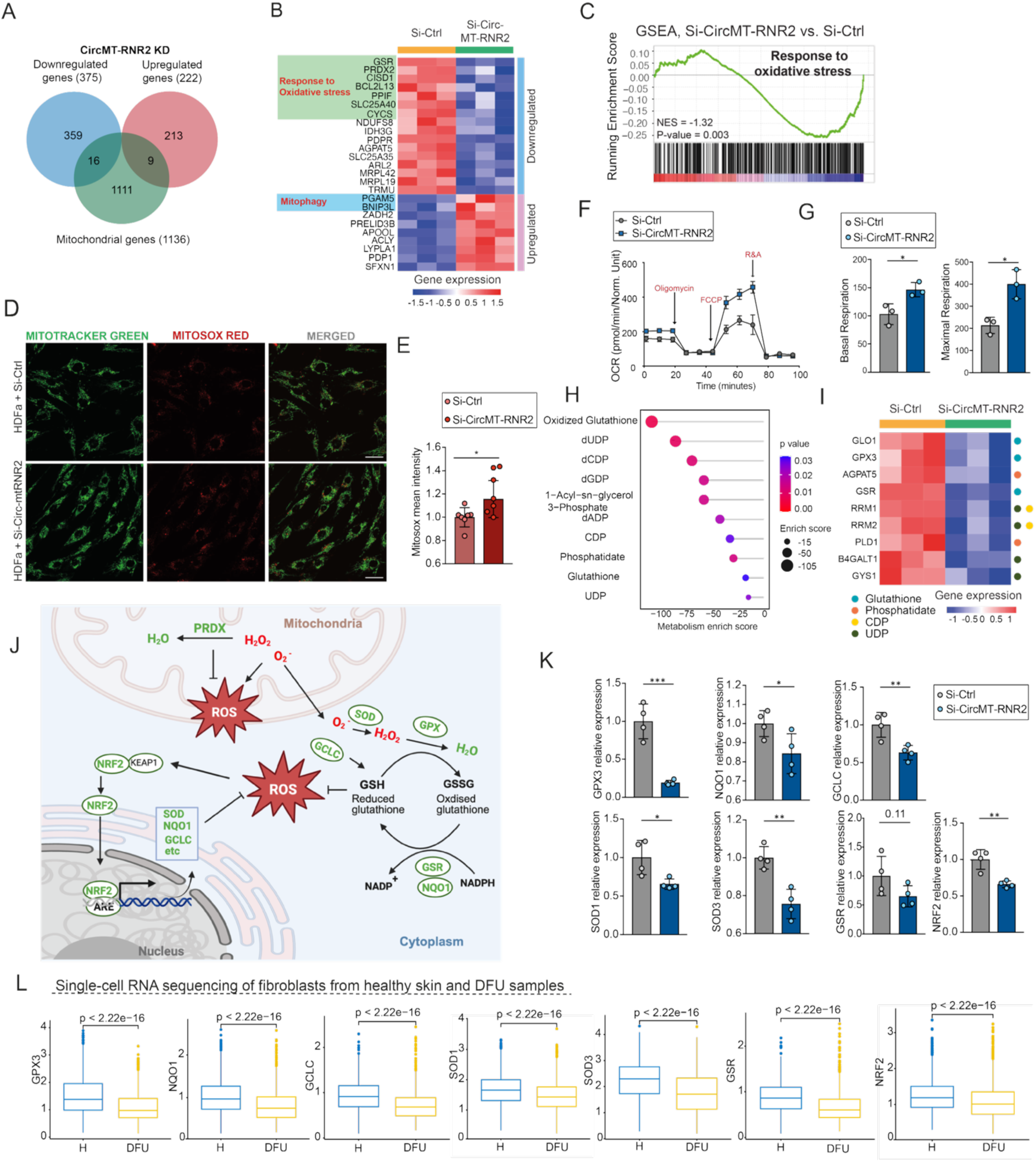
CircMT-RNR2 safeguards mitochondrial redox homeostasis. **(A)** Venn diagram showing overlap between differentially expressed genes in HDFa with circMT-RNR2 knockdown (KD) and mitochondrial genes. **(B)** Heatmap of up- and downregulated mitochondrial genes and their functions in HDFa with circMT-RNR2 KD. **(C)** Gene Set Enrichment Analysis (GSEA) of oxidative stress response–related genes following circMT-RNR2 KD. **(D)** Representative confocal images of HDFa cells stained with MitoTracker and MitoSOX after si-Ctrl or si-CircMT-RNR2 treatment; scale bar, 30 µm (n=7). **(E)** Quantification of MitoSOX mean fluorescence intensity. **(F-G)** Seahorse analysis of mitochondrial respiration showing oxygen consumption rate (OCR), basal, and maximal respiration in HDFa with circMT-RNR2 KD (n=3). **(H-I)** Metabolomic pathway analysis of genes downregulated upon circMT-RNR2 knockdown and corresponding heatmap of affected metabolic pathway–associated genes. **(J)** Schematic of antioxidant defense and ROS homeostasis. **(K)** qRT-PCR of antioxidant genes (GPX3, NQO1, GCLC, SOD1, SOD3, GSR, NRF2) in HDFa cells with circMT-RNR2 KD (n=4). **(L)** Single-cell RNA-sequencing analysis of the same antioxidant genes in dermal fibroblasts from healthy skin (H) (8425 cells from 9 donors) and DFU samples (10692 cells from 11 donors). *P < 0.05, **P < 0.01, ***P < 0.001 (unpaired Student t test, or Mann–Whitney U test).

To directly assess oxidative stress, we co-stained HDFa with MitoTracker™ Green (mitochondria) and MitoSOX Red (mitochondrial superoxide). CircMT-RNR2 knockdown increased mitochondrial ROS levels, indicating elevated oxidative stress (**Fig. 3D–E**). Similar ROS elevation was observed under hyperglycemic conditions, suggesting that circMT-RNR2 deficiency mimics hyperglycemia-induced stress (**Fig. S2A–B**). Seahorse analysis further showed that circMT-RNR2 knockdown increased oxygen consumption rate (OCR), as well as basal and maximal respiration (**Fig. 3F–G**). Interestingly, enhanced mitochondrial respiration coupled with elevated ROS has been reported in chronic diabetic complications, including DFU, where it impairs fibroblast function and wound healing (*38–40*).

Metabolic pathway analysis revealed that circMT-RNR2 silencing affected the oxidized glutathione pathway, indicating oxidative stress–related damage (**Fig. 3H**) (*41*). This was associated with loss of *GSR*, essential for glutathione recycling (*42*), and *GLO1*, which detoxifies harmful glycolysis byproducts (*43*) **(Fig. 3H–J)**. Reduced GLO1 activity promotes advanced glycation end-product (AGE) accumulation in diabetic models, a known driver of diabetic complications (*44*). Another key antioxidant gene, GPX3, was also downregulated; as a potent plasma peroxidase, GPX3 protects cells by reducing hydrogen peroxide and lipid hydroperoxides **(Fig. 3I–J)**(*45*). qRT-PCR confirmed reduced expression of *GPX3, GSR* and additional antioxidants, including *NRF2*, *NQO1, GCLC,* SOD1 and SOD3, upon circMT-RNR2 silencing (**Fig. 3K**). NRF2 is the master regulator of the antioxidant response, translocating to the nucleus under oxidative stress to induce enzymes such as NQO1 (quinone detoxification), GCLC (glutathione synthesis), and SOD1/3 (superoxide dismutation in cytoplasm and extracellular space), which together form a coordinated defense network that maintains redox balance and protects cells from oxidative damage (**Fig. 3J**)(*46–50*). Importantly, re-analysis of published single-cell RNA-sequencing data from healthy skin (n = 9) and DFU wound-edge tissue (n = 11) revealed consistent downregulation of these antioxidant genes in dermal fibroblasts from DFU compared with non-diabetic skin, underscoring the clinical relevance of circMT-RNR2 deficiency (**Fig. 3L**) (*51*).

CircMT-RNR2 silencing also upregulated *PGAM5* and *BNIP3L*, genes involved in mitophagy (*52–55*) (**Fig. 3B**). Western blotting confirmed increased levels of NIX (a mitophagy receptor) (*53*) and SIRT3 (a mitophagy activator) (*56*), alongside decreased TOMM20 (a mitochondrial integrity marker) (*57*), indicating enhanced mitophagy and mitochondrial damage (**Fig. S2C**). Analysis of global autophagic activity revealed that circMT-RNR2 knockdown increased LC3-I and LC3-II accumulation after ammonium chloride (NH₄Cl) treatment, which inhibits lysosomal degradation by neutralizing acidity and causes autophagosome accumulation (*58, 59*). This demonstrates elevated autophagic flux following circMT-RNR2 silencing (**Fig. S2D**). Collectively, circMT-RNR2 plays a critical role in maintaining mitochondrial redox balance by regulating antioxidant defense and oxidative stress response genes. Its loss increases oxidative stress, mitochondrial damage, and mitophagy, mimicking diabetic-related cellular dysfunction.

### CircMT-RNR2 interacts with and stabilizes PRDX3

To investigate the molecular mechanism of circMT-RNR2, we profiled its protein interactome using RNA pulldown in HDFa cells (**Fig. 4A**) (*60*). A biotin-labeled probe targeting the junction site of circMT-RNR2 enriched circMT-RNR2 over 4,000-fold compared with a control probe, as confirmed by qRT-PCR (**Fig. 4B**). Silver staining revealed more proteins pulled down by the circMT-RNR2 probe than by the control (**Fig. S3A**). Label-free mass spectrometry identified 37 proteins significantly enriched with the circMT-RNR2 probe (**Fig. 4C, Table S7**). Although the pulldown used whole-cell lysate, 5 of the top 11 enriched proteins were mitochondrial, supporting a mitochondrial localization and function for this circRNA. Among these, Peroxiredoxin 3 (PRDX3)—a mitochondria-specific ROS scavenger—drew particular attention given circMT-RNR2’s role in mitochondrial redox balance (*61*). RNA-Protein Interaction Prediction (RPISeq) analysis predicted a strong circMT-RNR2–PRDX3 interaction, with both RF and SVM classifiers scoring above the 0.5 binding threshold **(Fig. 4D)** (*62*). RNA immunoprecipitation (RIP) with PRDX3 antibody confirmed this interaction: circMT-RNR2 was significantly enriched in the PRDX3-IP compared with IgG control (**Fig. 4E**).

**Figure 4.**
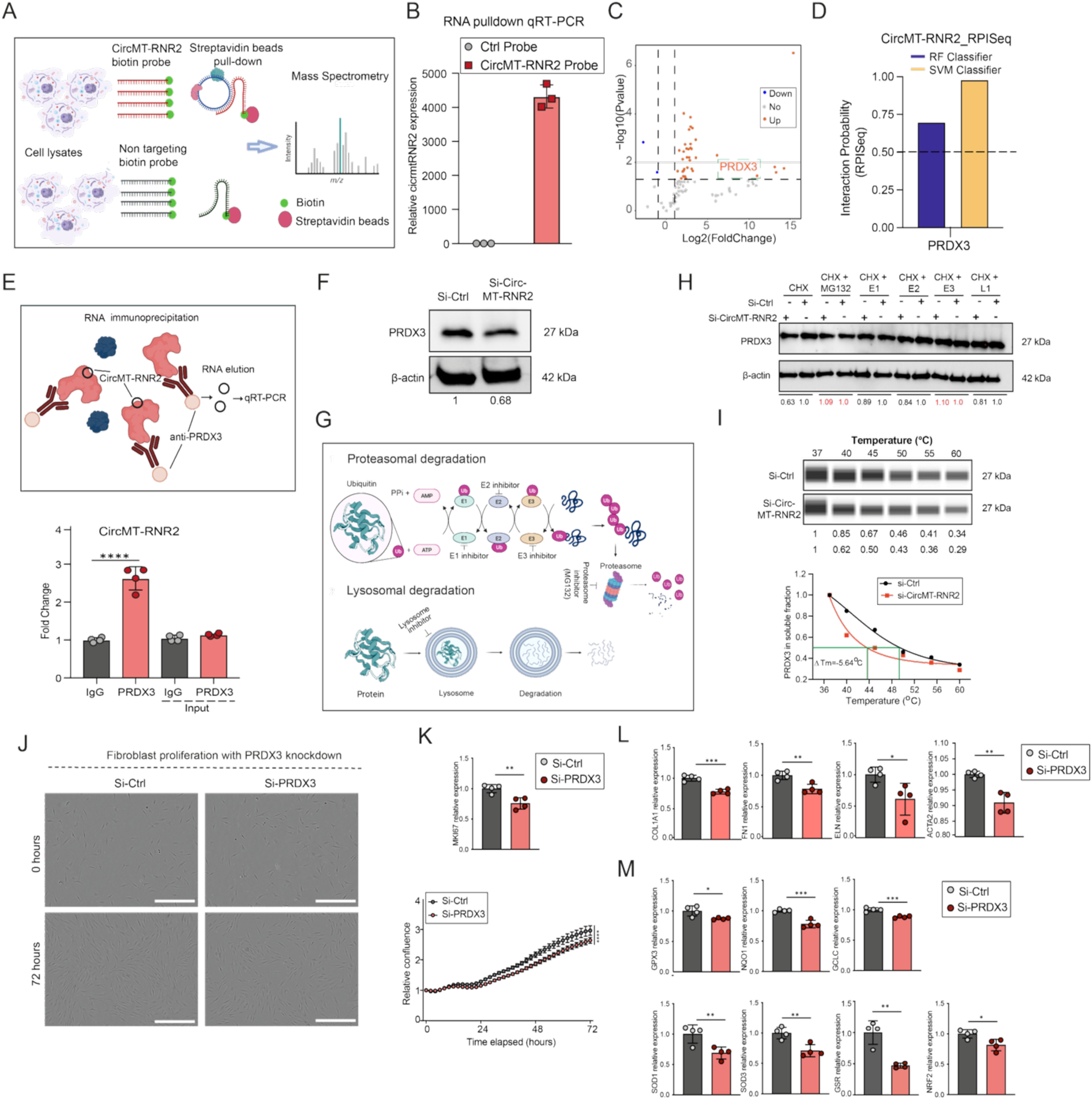
CircMT-RNR2 interacts with and stabilizes PRDX3. **(A)** Schematic of RNA pulldown using biotin-labeled circRNA probes and streptavidin beads followed by mass spectrometry (MS). **(B)** qRT-PCR validation of circMT-RNR2 enrichment after pulldown (n=3). **(C)** Volcano plot of proteins identified by MS. (**D**) RPISeq prediction showing strong interaction potential between PRDX3 and circMT-RNR2. **(E)** RNA immunoprecipitation (RIP) schematic (upper) and qRT-PCR detection of circMT-RNR2 in lysates immunoprecipitated with anti-PRDX3 or IgG control (n=4, lower). **(F)** Western blot of PRDX3 in HDFa cells with circMT-RNR2 knockdown; β-actin as control. **(G)** Schematic of protein degradation pathways. **(H)** Western blot of PRDX3 in circMT-RNR2–depleted fibroblasts treated with CHX alone or in combination with proteasomal (E1–E3) or lysosomal (L1) inhibitors; quantification relative to β-actin. **(I)** CETSA assay of PRDX3 stability: Simple Western (upper) and melting curves (lower) in HDFa cells with circMT-RNR2 silencing. **(J)** Proliferation assay of HDFa cells with PRDX3 knockdown and quantification of cell confluence (72 h, n=6); scale bar, 400 µm. **(K)** qRT-PCR of *MKI67* after PRDX3 knockdown (n=4). **(L)** qRT-PCR of *COL1A1, FN1, ELN*, and *ACTA2* in HDFa cells with PRDX3 knockdown (n=4). **(M)** qRT-PCR of antioxidant genes (GPX3, NQO1, GCLC, SOD1, SOD3, GSR, NRF2) after PRDX3 knockdown (n=4). *P<0.05, **P<0.01, ***P<0.001, ****P<0.0001 (unpaired Student t.test, or two-way ANOVA and multiple comparisons).

Furthermore, we found that PRDX3 protein levels decreased upon circMT-RNR2 knockdown (**Fig. 4F**). To assess whether circMT-RNR2 regulates PRDX3 production or degradation, fibroblasts were treated with cycloheximide (CHX) to block translation, in combination with lysosomal or proteasomal inhibitors to prevent protein degradation (**Fig. 4G**) (*63*). Blocking degradation—but not translation—restored PRDX3 levels in circMT-RNR2-depleted cells, suggesting that circMT-RNR2 stabilizes PRDX3 protein (**Fig. 4H**). This was supported by Cellular Thermal Shift Assay (CETSA), where circMT-RNR2 silencing reduced PRDX3 melting temperature, indicating decreased protein stability (**Fig. 4I**) (*64*).

PRDX3 is essential for mitochondrial function and oxidative stress protection (*61*). Its knockdown increases mitochondrial ROS, disrupts mitochondrial function, and impairs cancer cell proliferation (*65–67*). In fibroblasts, we showed that PRDX3 silencing (**Fig. S3B**) reduced cell proliferation and *MKI67* expression – a proliferation marker (**Fig. 4J, K**), decreased ECM gene expression (*COL1A1, FN1, ELN*), and lowered *ACTA2* levels (**Fig. 4L**). It also reduced the expression of antioxidant enzymes (*GPX3, NQO1, GCLC, SOD1*, *SOD3, GSR*) and the master antioxidant regulator *NRF2* (**Fig. 4M**). Thus, PRDX3 knockdown phenocopied circMT-RNR2 depletion, impairing fibroblast proliferation, ECM production, contractility, and mitochondrial antioxidant defense. These results support a model in which circMT-RNR2 binds and stabilizes PRDX3 to preserve mitochondrial redox homeostasis.

### CircMT-RNR2 is essential for wound healing in murine *in vivo* wound model

Given that circMT-RNR2 is conserved between humans and mice (**Fig. 1D**), we examined its expression in a murine wound healing model. CircMT-RNR2 was upregulated in wound fibroblasts compared with skin fibroblasts and was more abundant in dermis than epidermis, mirroring its distribution in human skin (**Fig. 5A, B**). In wild-type mice, circMT-RNR2 levels increased progressively during wound healing, whereas in db/db mice—a type 2 diabetes model—its expression remained unchanged, consistent with reduced circMT-RNR2 in human DFU (**Fig. 5C, Fig. S4A, B**) (*68*).

**Figure 5.**
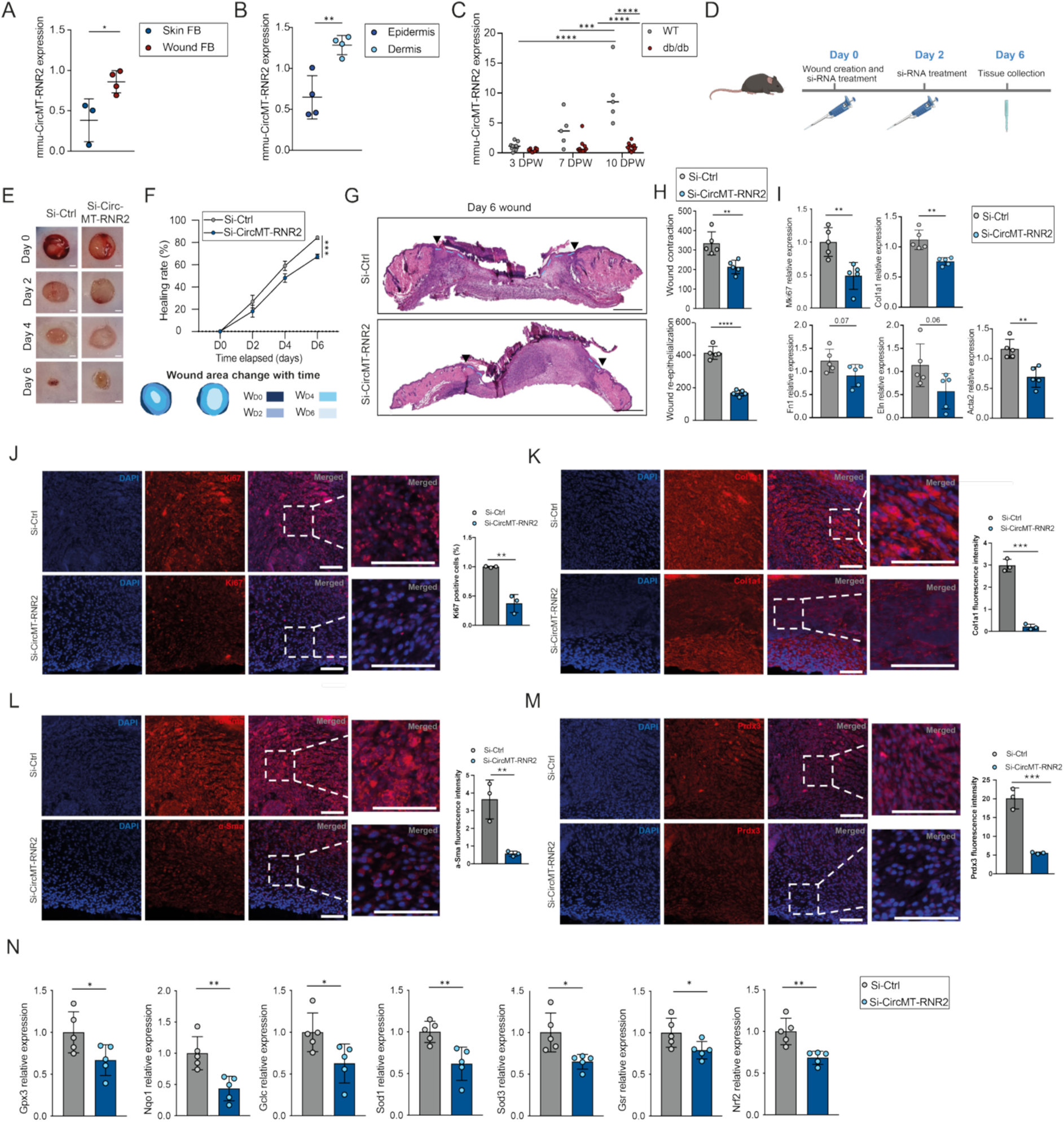
CircMT-RNR2 is required for wound closure *in vivo*. qRT-PCR analysis of mmu_circMT-RNR2 in **(A)** mouse skin (n=3) and wound fibroblasts (n=4); **(B)** murine epidermis vs. dermis (n=4); **(C)** during wound healing in wild-type and db/b mice (days post-wounding, DPW; n=5-10). **(D)** Schematic of topical siRNA treatment targeting mmu_circMT-RNR2. **(E)** Macroscopic wound images (Day 0–Day 6) after siRNA treatment (n=5). Scale bar = 1000 µm. **(F)** Quantification of wound healing rate (%) from **(E). (G)** Representative H&E images of D6 wounds after Si-Ctrl or Si-mmu-CircMT-RNR2 treatment (n=5). Scale bar = 500 µm. Dashed blue lines mark new epidermis; arrowheads indicate initial wound edges. **(H)** Quantification of re-epithelialization and contraction from **(G). (I)** qRT-PCR of *Mki67*, *Col1a1*, *Fn1*, *Eln* and *Acta2* from D6 wound biopsies (n=5). (**J–M**) Representative immunofluorescence images of mouse wounds stained for (**J**) Ki67, (**K**) Col1a1, (**L**) α-Sma, and (**M**) Prdx3, with signal quantification (n=3). Scale bar = 100 µm. **(N)** qRT-PCR of *Gpx3*, *Nqo1*, *Gclc*, *Sod1*, *Sod*3, *Gsr* and *Nrf2* in mouse dermis after siRNA treatment (n=5). *P<0.05, **P<0.01, ***P<0.001, ****P<0.0001 (unpaired t-test, or one-way ANOVA and Turkey’s multiple comparison test, or two-way ANOVA and multiple comparisons).

To test whether circMT-RNR2 is required for wound repair, we topically applied si-CircMT-RNR2 or si-Ctrl to dorsal skin wounds immediately after injury and again two days later, monitoring healing over six days (**Fig. 5D**). Si-CircMT-RNR2 treatment significantly reduced dermal circMT-RNR2 expression through day 6 (**Fig. S4C**) without affecting body weight, indicating no systemic toxicity (**Fig. S4D**). Macroscopically, circMT-RNR2 knockdown delayed wound closure (**Fig. 5E, F**). Hematoxylin and Eosin (H&E) staining of day 6 wounds confirmed reduced contraction and re-epithelialization (**Fig. 5G, H**). qRT-PCR revealed lower expression of the proliferation marker *Mki67*, ECM genes (*Col1a1*, *Fn1*, *Eln*), and the contractility gene *Acta2* (**Fig. 5I**). Reduced *Mki67*, *Col1a1*, and *Acta2* protein levels were confirmed by immunofluorescence (**Fig. 5J–L**). Notably, circMT-RNR2 knockdown markedly decreased Prdx3 protein in wound tissue (**Fig. 5M**). In parallel, it suppressed the expression of antioxidant enzymes (*Gpx3*, *Nqo1*, *Gclc*, *Sod1*, *Sod3, Gsr*) and the master antioxidant regulator *Nrf2* (**Fig. 5N**).

Collectively, these results demonstrate that circMT-RNR2 is critical for wound healing *in vivo*, acting through Prdx3 stabilization and the preservation of mitochondrial redox balance, thereby protecting cells from oxidative stress–induced damage.

### CircMT-RNR2 promotes healing of human *ex vivo* wounds

To evaluate the translational relevance of our findings, we investigated the role of circMT-RNR2 in wound repair using a human *ex vivo* skin model (**Fig. 6A**). In this system, wounds are introduced into surgically discarded human skin samples, which are then maintained under standard culture conditions. This approach preserves native tissue architecture, allowing the study of human skin repair in an environment that closely resembles *in vivo* conditions (*69, 70*).

**Figure 6:**
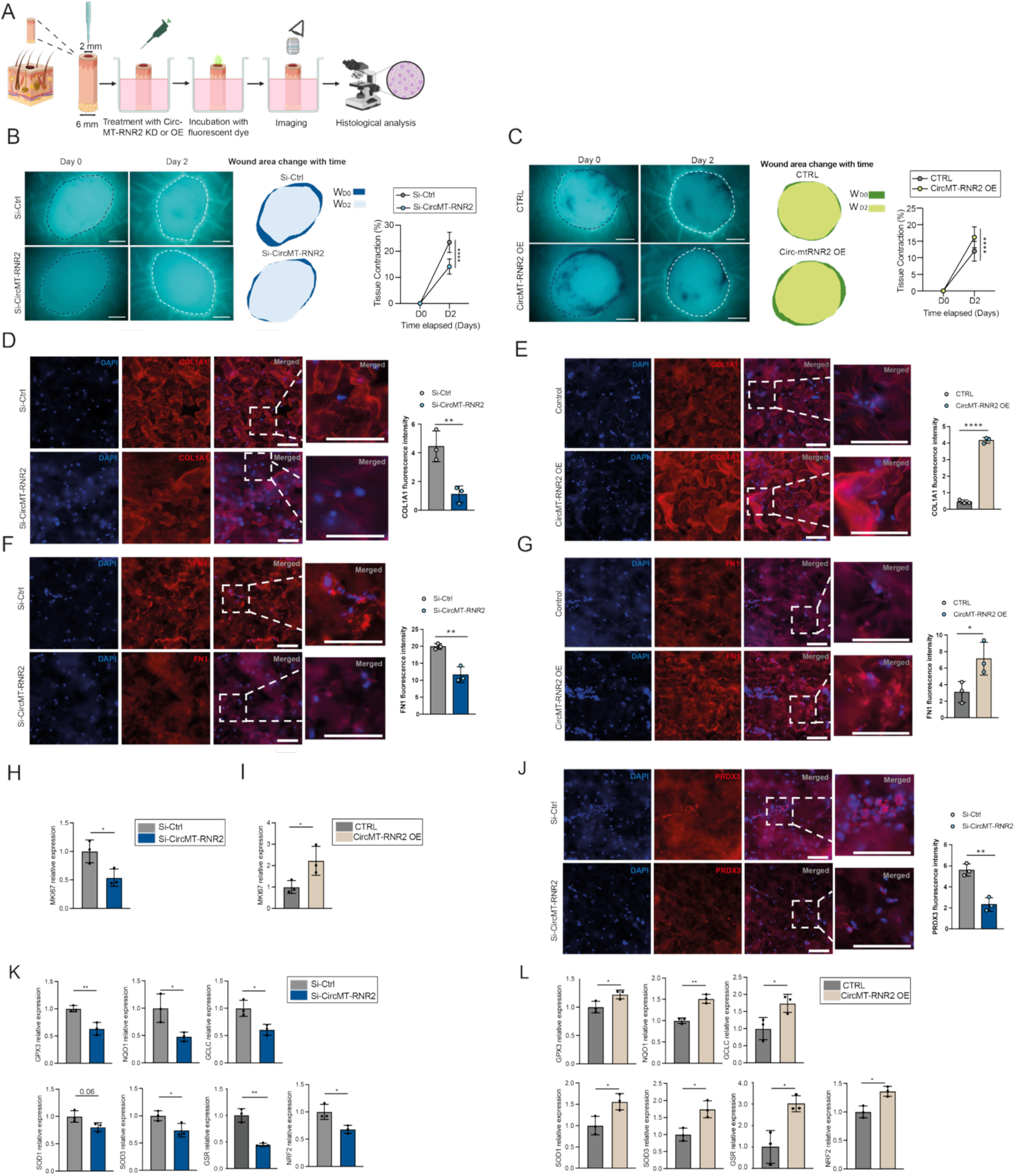
CircMT-RNR2 promotes healing of human *ex vivo* wounds. **(A)** Schematic illustration of topical treatment of human *ex vivo* wounds with si-RNA targeting circMT-RNR2 (KD) or circMT-RNR2 overexpression vector (OE). **(B-C)** Fluorescent images of wound areas at D0 and D2. Wounds treated with Si-Ctrl or Si-CircMT-RNR2 **(B)** and Control or circMT-RNR2 OE **(C)** are shown (left); wound edges are indicated by dashed lines. Areas within the wound edge (IW) at each time point are illustrated (right). Scale bars: 500 µm. Wound contraction (%) was calculated as ΔIWD2 /IWD0×100% (n=3/group). **(D-G)** Representative immunofluorescence images showing COL1A1 (**D, E**) and FN1 (**F, G**) in wounds with circMT-RNR2 KD (**D, F**) or OE (**E, G**) for 7 days. Scale bars: 100 µm (**H**-**I**) qRT-PCR analysis of *MKI67* in these treated human wounds. **(J)** Immunofluorescence images of PRDX3 in wounds with circMT-RNR2 KD. (**K**-**L**) qRT-PCR analysis of oxidative stress-related genes (*GPX3*, *NQO1*, *GCLC*, *SOD1*, *SOD3*, *GSR*, *NRF2)* in treated dermis (n=3). *P<0.05, **P<0.01, ***P<0.001, ****P<0.0001 (unpaired Student t-test, or two-way ANOVA and multiple comparisons).

We modulated circMT-RNR2 levels in *ex vivo* wounds by silencing with si-CircMT-RNR2 or by overexpressing (OE) using a plasmid based on a previously established method for mitochondrial circRNA expression (*26*) (**Fig. S5A**). qRT-PCR confirmed efficient mitochondrial overexpression of circMT-RNR2 in human dermal fibroblasts without affecting MT-RNR2 expression (**Fig. S5B-D**), and successful circMT-RNR2 overexpression in the wound dermis of the *ex vivo* model (**Fig. S5E**).

Functionally, silencing circMT-RNR2 impaired tissue contraction, whereas overexpression promoted it (**Fig. 6B, C**). Immunofluorescence analysis revealed that knockdown of circMT-RNR2 decreased dermal expression of COL1A1 and FN1, whereas overexpression increased their levels (**Fig. 6D–G**). Similarly, silencing circMT-RNR2 reduced, while overexpression increased, the proliferation marker *MKI67* (**Fig. 6H, I**). Notably, PRDX3 protein was reduced in circMT-RNR2–silenced wounds, supporting circMT-RNR2’s role in stabilizing PRDX3 during human wound repair (**Fig. 6J**). Moreover, qRT-PCR revealed that circMT-RNR2 knockdown suppressed antioxidant enzymes (*GPX3, NQO1, GCLC, SOD1*, *SOD3, GSR*) and the master antioxidant regulator *NRF2,* while overexpression enhanced their expression (**Fig. 6K, L**). Collectively, these results highlight circMT-RNR2 as a critical regulator of skin repair and a potential therapeutic target.

## DISCUSSION

Mitochondria-encoded circular RNAs (mecciRNAs) are a newly recognized class of non-coding RNAs that expand our understanding of how mitochondria regulate cellular homeostasis and stress adaptation (*71*). Although hundreds of mecciRNAs have been cataloged in human and murine cells, only a few have been functionally characterized, each revealing important roles in mitochondrial biology (*11–13, 15*). Here, we identify circMT-RNR2 as a novel functional mecciRNA essential for wound repair. By stabilizing the antioxidant protein PRDX3, circMT-RNR2 safeguards mitochondrial redox homeostasis, thereby supporting fibroblast activity crucial for tissue repair. Its deficiency in DFUs compromises fibroblast function and delays healing, providing the first evidence that a pathophysiologically relevant mecciRNA contributes to human tissue repair (**Fig. 7**).

**Figure 7.**
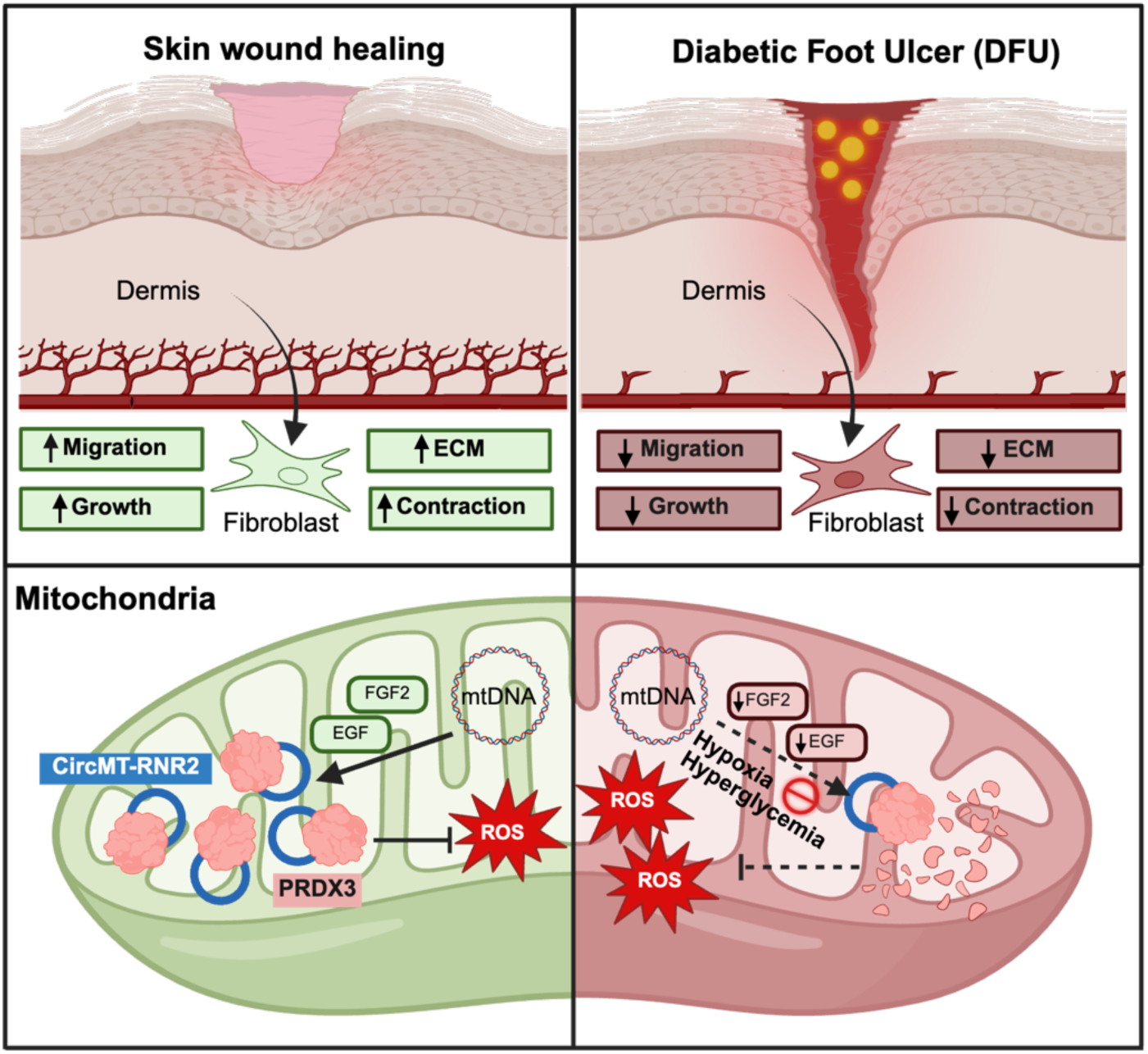
Summary of study findings. CircMT-RNR2 promotes wound healing by enhancing fibroblast proliferation, migration, contraction, and extracellular matrix (ECM) production. It sustains mitochondrial redox balance by stabilizing the antioxidant protein PRDX3, thereby reducing ROS-induced damage. In diabetic foot ulcers, hypoxia, hyperglycemia, and reduced FGF2 and EGF expression lower circMT-RNR2 levels, impairing fibroblast function and delaying healing.

Mitochondrial transcripts are synthesized as long polycistronic RNAs, then processed into mRNAs, tRNAs, and rRNAs primarily through tRNA-guided cleavage by RNase P and RNase Z (*72–74*). While the tRNA punctuation model explains much of this process, the existence of non–tRNA-flanked transcripts and newly identified mitochondrial non-coding RNAs suggests alternative RNA processing pathways (*15*). Unlike nuclear circRNAs, which form through backsplicing, mecciRNAs are unlikely to use canonical splicing machinery, which is absent in mitochondria (*11, 75*). Notably, circMT-RNR2 and many other mecciRNAs harbor conserved repetitive sequences at their junctions that may facilitate mitochondrial RNA ligation (**Fig. 1C**) (*11, 13*). Although nuclear mitochondrial DNA segments could raise questions about circMT-RNR2’s genomic origin, our data show that inhibiting mitochondrial RNA polymerase reduces circMT-RNR2 levels without affecting nuclear RNAs, supporting its mitochondrial origin (**Fig. 1D**) (*76*). These observations suggest that mecciRNAs arise via a unique and currently uncharacterized mitochondrial mechanism, warranting further investigation.

Our data show that circMT-RNR2 expression in human dermal fibroblasts is dynamically regulated—upregulated by growth factors such as EGF and FGF2, but suppressed under hyperglycemic or hypoxic conditions. This regulation mirrors that of nuclear circRNAs, indicating that circMT-RNR2 is not transcriptional byproduct but a responsive regulatory molecule. In DFUs, loss of proliferating fibroblasts correlates with reduced EGF signaling (*77*), and diminished FGF2 further impairs fibroblast activation and angiogenesis (*78*). Combined with the hostile metabolic and hypoxic microenvironment of DFUs, these deficiencies likely converge to suppress circMT-RNR2 expression in chronic wounds.

Redox homeostasis is crucial for wound healing, and our findings show that circMT-RNR2 supports mitochondrial ROS balance by strengthening the antioxidant defense system. We identify PRDX3, a mitochondria-specific ROS scavenger that is induced under oxidative stress (*61, 79*), as a direct binding partner of circMT-RNR2. This interaction prevents PRDX3 degradation, thus limiting mitochondrial ROS accumulation. Given that excessive ROS induces FOXM1 expression (*80*) and that FOXM1 transcriptionally regulates PRDX3 (*65*), circMT-RNR2 may reinforce PRDX3 levels through both transcriptional activation (likely via FOXM1) and protein stabilization. This dual regulation highlights the pivotal role of circMT-RNR2 in supporting mitochondrial redox homeostasis, thereby preserving fibroblast viability under oxidative stress, which may be especially critical in the chronically inflamed, oxidative environment of DFUs.

In summary, this study identifies circMT-RNR2 as a novel regulator of mitochondrial antioxidant defense, with direct relevance to tissue repair. Beyond establishing the first functional link between a mecciRNA and human wound healing, our findings broaden the understanding of mitochondrial RNA biology and highlight circMT-RNR2 as a potential therapeutic target for chronic, non-healing wounds in diabetes.

## MATERIALS AND METHODS

### Human skin specimens

Human skin samples and DFU tissues were collected at the Second Hospital of Dalian Medical University, China (**Table S1**). DFU patients enrolled in the study had non-healing ulcers persisting for >2 months despite conventional treatment. Tissue was obtained from the non-healing wound edges using a 4-mm biopsy punch. Human skin for cell isolation and *ex vivo* wound model experiments was sourced from discarded abdominal tissue following plastic surgery at the Department of Reconstructive Plastic Surgery, Karolinska University Hospital, Sweden (**Table S1**). All participants provided written informed consent. The study protocols were approved by the Ethics Committee of The Second Hospital of Dalian Medical University (2015–102, 2016–28) and the Stockholm Regional Ethics Committee (2019–02335), and conducted in accordance with the Declaration of Helsinki.

### Microarray analysis

Transcriptomic profiling was performed by employing the Affymetrix Human Clariom™ S Assay (Thermo Fisher Scientific) at the Bioinformatics and Expression Analysis (BEA) Core Facility, Karolinska Institutet. RNA quality and quantity were assessed using a NanoDrop 1000 spectrophotometer (Thermo Fisher Scientific) and an Agilent 2200 TapeStation with RNA ScreenTape (Agilent, Santa Clara, CA). A total of 150 ng RNA was used for cDNA synthesis according to the GeneChip WT PLUS Reagent Kit protocol (Thermo Fisher Scientific). Standard Affymetrix procedures, including hybridization, fluidics processing, and scanning, were followed. Genes exhibiting |log2FoldChange| >1.5 with FDR < 0.05 were considered differentially expressed (DEGs). Results were visualized as heatmaps and volcano plots generated using the “heatmap” and “ggplot2” R packages. Overlaps between DEGs and mitochondrial genes were illustrated with a Venn diagram. The dataset is available at the Gene Expression Omnibus under accession number GSE301636.

To investigate the biological pathways associated with DEGs, Kyoto Encyclopedia of Genes and Genomes (KEGG) pathway enrichment analysis was conducted using the R package clusterProfiler, applying a significance cutoff of p < 0.05. Additionally, Gene Set Enrichment Analysis (GSEA) was performed to compare functional differences between groups. Metabolomic enrichment and transcription factor prediction analyses were carried out using the Enrichr platform (https://maayanlab.cloud/Enrichr/)(81). Lollipop plots depicting differences in metabolomic pathways and transcription factors between HDFa si-Ctrl and HDFa si-CircMT-RNR2 groups were generated using the R package ggplot2.

### Cell culture and and treatments

Primary human adult dermal fibroblasts (HDFa; Cascade Biologics, Portland, OR) were cultured in Dulbecco’s Modified Eagle Medium high glucose (DMEM, ThermoFisher Scientific) supplemented with 10% heat inactivated Fetal Bovine Serum (HI-FBS) and 1% antibiotic coktail (penicillin/ streptomycin) at 37°C in 5% CO2. To mimic hypoxia, cells were incubated under hypoxic conditions within a chamber set to 5% oxygen (O₂).

To investigate factors potentially regulating circMT-RNR2 expression, HDFa cells were treated with the following cytokines and growth factors: IL-1α (20 ng/ml), IL-6 (50 ng/ml), IL-8 (50 ng/ml), IL-22 (30 ng/ml), IL-36α (100 ng/ml), TNF-α (50 ng/ml), TGF-β1 (20 ng/ml), TGF-β2 (10 ng/ml), TGF-β3 (20 ng/ml), BMP-2 (100 ng/ml), EGF (20 ng/ml), IGF-1 (20 ng/ml), FGF-2 (30 ng/ml), VEGFA (20 ng/ml), HB-EGF (20 ng/ml) or PBS as a control. Treatment was applied for 24 hours, after which circMT-RNR2 expression was analyzed by qRT-PCR. All cytokines and growth factors were obtained from ImmunoTools (Friesoythe, Germany) or R&D Systems (Minneapolis, MN).

To inhibit mitochondrial transcription, cells were treated with 5 μM or 10 μM IMT1 (HY-134539, MedChemExpress) for 200 hours to block mitochondrial RNA polymerase. The culture medium was replaced every other day with fresh medium containing the same concentration of IMT1.

To assess autophagy, HDFa cells transfected with Si-Ctrl or Si-CircMT-RNR2 were treated with 50 nM ammonium chloride (NH₄Cl; HY-Y1269C, MedChemExpress) for 3 hours. Proteins were then extracted and analyzed by Western blot.

To study the functions of circMT-RNR2/PRDX3 in fibroblasts, knockdown and overexpression experiments were performed. For knockdown, cells at 70% confluence were transfected with predesigned siRNA targeting circMT-RNR2 (Si-CircMT-RNR2, Dharmacon) or with siRNA targeting PRDX3 (Si-PRDX3, Dharmacon) and a non-targeting control siRNA (Si-Ctrl, Dharmacon) for 24 hours using RNAiMAX as transfection reagent, in nutrient depleted, antibiotic free medium. To overexpress circMT-RNR2, fibroblasts at 80–90% confluence were transfected with circMT-RNR2 overexpression plasmid using Lipofectamine™ 3000 for 24 hours.

To assess endogenous PRDX3 protein turnover, HDFa cells were treated for 2 hours with 0.5 μM MG132 (Sigma-Aldrich, Cat. No. M7449), 5 μg/mL cycloheximide (CHX, Sigma-Aldrich, Cat. No. C4859), 10 μM ubiquitin E1 inhibitor PYR-41 (Sigma-Aldrich, Cat. No. 662105), 2 μM UbcH13 E2 inhibitor (Sigma-Aldrich, Cat. No. 662107), 10 μM Heclin E3 inhibitor (Sigma-Aldrich, Cat. No. SML1396), and 1 μM Bafilomycin L1 (Sigma-Aldrich, Cat. No. 88899-55-2). Following treatment, cells were washed with PBS, and proteins were isolated for Western blot analysis.

### Proliferation and migration assays

HDFa cell proliferation and migration were assessed using the IncuCyte live-cell imaging system (Sartorius, Germany). For the proliferation assay, cells were seeded at low density (typically 20,000 cells/well) in 12-well plates (Sarstedt, Germany) and allowed to adhere overnight. For the migration assay, ImageLock 96-well plates (Essen Bioscience, Ann Arbor, MI) were pre-coated with Collagen I and incubated overnight. HDFa cells were seeded at 15,000 cells per well and allowed to adhere for several hours or overnight. Once a confluent monolayer was formed, a scratch was created using the IncuCyte wound maker (Essen Bioscience). Cells were then imaged every two hours, and migration was quantified with the IncuCyte ZOOM software. Proliferation marker *MKI67* was measured by qRT-PCR.

### Mitotracker green and Mitosox red staining

After incubation in either high- or low-glucose medium and exposure to hypoxic conditions or following transfection, cells were stained with MitoSOX Red (Thermo Fisher Scientific), a mitochondrial superoxide indicator, at 500 nM for 30 minutes according to the manufacturer’s instructions. Subsequently, MitoTracker Green (Thermo Fisher Scientific) was added at 100 nM for 10 minutes. Cells were then washed with HBSS and imaged using a ZEISS confocal microscope at the Biomedicum Imaging Core (BIC), Karolinska Institutet. Fluorescence intensity was quantified using ImageJ.

### Analysis of single-cell RNA-sequencing data

We analyzed publicly available scRNA-seq data of diabetic foot ulcers (DFUs) from Theocharidis *et al.* (GSE165816) (*51*), including samples from 9 healthy controls and 11 DFU patients. Cells with <500 detected genes, >20% mitochondrial content, or <1,000 total counts were removed. After quality control, we applied our previously described workflow (*77*), including normalization, identification of highly variable features, batch correction, dimensionality reduction, and unsupervised clustering (resolution 0.8). Cell type annotation was performed based on canonical marker gene expression as previously described (*51*) and gene expression analyses focused on fibroblast populations.

### RNA pulldown and mass spectrometry

The circMT-RNR2 pulldown assay was conducted using biotin-labeled probes and streptavidin-coated magnetic beads. A probe specific to circMT-RNR2 was designed, with a control probe included to assess specificity. Pulldown assay was carried out with the Pierce Co-Immunoprecipitation (Co-IP) Kit (Thermo Fisher Scientific) according to the manufacturer’s instructions. The captured RNA-protein complexes were eluted and bound proteins were analyzed by mass spectrometry at the Proteomics Biomedicum Core Facility (Karolinska Institutet), The mass spectrometry proteomics data have been deposited to the ProteomeXchange Consortium via the PRIDE (*82*) partner repository with the dataset identifier PXD067605 (Token: htLz3AQaMKxK). The associated RNAs were extracted using TRIzol and analyzed by qRT-PCR to confirm enrichment of circMT-RNR2.

### RNA immunoprecipitation

RNA immunoprecipitation (RIP) was performed using the Magna RIP RNA-Binding Protein Immunoprecipitation Kit (Millipore, Burlington, MA). HDFa cells were lysed in RIP lysis buffer, and 100 µL of the whole-cell extract was incubated with an anti-human PRDX3 antibody (ab222807, Abcam) conjugated to Protein A + G magnetic beads (Millipore) in RIP buffer. Normal rabbit IgG (Millipore) was used as a negative control. After incubation, proteinase K treatment was applied to digest proteins, and the immunoprecipitated RNA was isolated. The levels of circMT-RNR2 were then quantified by qRT-PCR.

### Mouse skin specimens and in vivo wound model

Mouse skin samples were collected from Eleven-week-old male C57BL/6J (wild-type) mice or diabetic db/db [BKS(D)-Lepr^db^/JOrlRj] mice on the C57BL/6J background (Janvier Labs, Le Genest-Saint-Isle, France) as we previously reported (*83*). Briefly, each mouse received 4-mm excisional wounds, and the excised skin served as an intact skin control. Wound-edge tissues were collected using a 6-mm punch at 3, 7, and 10 days post-wounding (DPW). The samples were then were used for RNA extraction and qRT-PCR analysis. For dermis and epidermis separation, samples were then washed twice with PBS to remove debris and incubated overnight at 4 °C with Dispase II (5 U/mL; Roche) to separate the epidermis from the dermis. The whole skin as well as the sperated dermis and epidermis were used for RNA extraction and qRT-PCR analysis. For cell isolation, the dermal tissue was minced into small fragments before enzymatic digestion at 37 °C for 3–4 h using the Whole Skin Dissociation Kit (Miltenyi Biotec, 130-101-540). The resulting suspension was filtered through a 70-μm mesh and centrifuged to obtain single cells. Fibroblasts were purified using the PDGFRα MicroBead Kit (Miltenyi Biotec, 130-101-502) following the manufacturer’s protocol, and the isolated cells were used for RNA extraction and qRT-PCR analysis.

An *in vivo* wound healing model was established using 8–9 weeks old male mice with C57BL/6 J background over a seven-day period. The dorsal area of each mouse was shaved and treated with depilatory cream to remove hair. Two 4 mm full-thickness wounds were created on the dorsal surface of each mouse. A total of 5 µL of treatment, containing either 0.1 nmol si-CircMT-RNR2 (n = 5) or si-Control (n = 5), along with in vivo-jetPEI (Cat. 201-10 G, Polyplus-transfection, France), were applied topically at two time points: Day 0 and Day 2. The wounds were covered with Tegaderm film dressings. Pain relief was provided via intramuscular administration of Temgesic (0.003 mg/mL) at a dose of 100 µL per 10 g of body weight on Day 0 and Day 1. Wound images were captured on Days 0, 2, 4, and 6 to document the healing process. On Day 6, mice were euthanized in a CO₂ chamber, and wound tissues were harvested using a 6 mm biopsy punch. The collected biopsies were used for RNA extraction and qRT-PCR analysis to assess gene expression, as well as for histological examination via H&E staining.

### Human ex vivo wound model

Skin from discarded surgical procedures was cleaned with 70% ethanol, and partial-thickness wounds (restricted to the dermis) were created using a 2 mm biopsy punch. These wounds were then excised with a 6 mm punch and placed in 8-well chamber plates for *in vitro* culture. Samples were maintained in DMEM supplemented with 10% FBS, 1% penicillin– streptomycin, and 1% antifungal agent (Thermo Fisher Scientific). 80 µL of medium was added to each biopsy, ensuring the epidermis remained exposed to air to preserve a liquid–air interface. Cultures were incubated at 37 °C in a humidified 5% CO₂ atmosphere.

For treatments, si-CircMT-RNR2 or si-Ctrl (n = 3 per condition) was applied topically at 1.5 µg/µL, mixed with 10% glucose, PBS, and JetPei transfection reagent (Polyplus-transfection). CircMT-RNR2 overexpression (OE) and control plasmids were applied at 3 µg/µL. A total of 10 µL treatment was administered on Days 0 and 2. Wound samples were collected six days post-injury for RNA extraction and qRT-PCR to evaluate transfection efficiency and gene expression. Wound closure was assessed using CellTracker™ Green CMFDA Dye (Invitrogen). Briefly, 4 µL dye (50 µM) was added to each tissue and incubated at 37 °C with 5% CO₂ for 30 minutes. After PBS washing, wounds were imaged with a Nikon Eclipse Ni-E fluorescence microscope, and wound areas were quantified using ImageJ.

### Statistics

The number of biological replicates for each experiment is specified in the respective methods sections and figure legends. Comparisons between two groups were made using either a paired or unpaired Student’s t-test, while comparisons involving multiple groups were analyzed using one way or two-way analysis of variance (ANOVA), comparisons between single cell samples using Mann–Whitney U test. Statistical significance was determined at a threshold of P < 0.05. Statistical analyses were conducted using GraphPad Prism software version 9 and R software version 4.3.3.

## Acknowledgments and funding

We are deeply grateful to the tissue donors for their participation. We thank the Bioinformatics and Expression Analysis (BEA) core facility, supported by the Board of Research at Karolinska Institutet, for microarray analysis, as well as the Proteomics Biomedicum Core Facility, the Histology Core Facility, and the Biomedicum Imaging Core (BIC) for their support. This study was funded by the Swedish Research Council (2020-01400, 2024-02739 to NXL), Welander and Finsens Foundation (Hudfonden) (NXL, GN), Åke Wibergs Stiftelse (GN), Ming Wai Lau Centre for Reparative Medicine (NXL), LEO Foundation (LF-AW_EMEA-20-400022, NXL), and a Postdoctoral Fellowship from the Strategic Research Programme in Diabetes at Karolinska Institutet (GN).

## Author contributions

NXL, GN, JG, AW, PS, and XZ conceived the study and designed the experiments. GN, JG and YX collected clinical samples and performed the experiments. GN and JG analyzed the data and prepared the figures. YC and ZL conducted bioinformatic analysis. NXL, GN and JG collaboratively prepared the original manuscript draft with the help from all authors.

## Materials and correspondence

Further information and requests for resources and reagents should be directed to and will be fulfilled by the lead contact, Ning Xu Landén.

