## Supplementary figures for "Mitochondrial CircRNA CircMT-RNR2 Safeguards Antioxidant Defense to Support Fibroblast Functions in Wound Repair"

**The PDF file includes:**

Materials and Methods

Table legends; Figures S1-S5

Table S1. Human sample information.

Table S2. DEG in HDFa with circMT-RNR2 knockdown.

Table S3. KEGG pathway analysis of DEG in HDFa with circMT-RNR2 knockdown.

Table S4. Transcription factor enrichment analysis of downregulated genes in HDFa with circMT-RNR2 knockdown.

Table S5. Table S5 FOXM1 regulated genes in in HDFa with circMT-RNR2 knockdown.

Table S6 Metabolic pathway analysis of DEG in HDFa with circMT-RNR2 knockdown.

Table S7. Mass spectrometry analysis of protein interactome of circMT-RNR2.

Table S8. List of reagents used in this study.

### MATERIALS AND METHODS

#### *Dermis and epidermis separation and cell isolation*

The skin samples were washed in PBS supplemented with 1% antifungal agent and 1% antibiotic cocktail (penicillin/streptomycin) (Thermo Fisher Scientific, Germany). The biopsies were then incubated in Dispase (5 U/mL, 370 mg) (Roche, Switzerland) for 12–16 hours, followed by a second wash with PBS containing 1% antifungal agent and 1% antibiotic cocktail. The epidermis was carefully separated from the dermis using forceps and stored at −80 °C until further analysis. For cell isolation, magnetic-activated cell sorting (MACS) was used to isolate keratinocytes (CD45- epidermal cells) and fibroblasts (CD90+ dermal fibroblasts) from the epidermis and dermis, following our previously described method(*1*).

#### *RNase R digestion, gel electrophoresis, and Sanger sequencing*

To verify the circular structure of circMT-RNR2, total RNA from human dermal fibroblasts was incubated with RNase R at 37 °C for 30 or 60 min, followed by heat inactivation at 70 °C for 10 min. Divergent primers spanning the junction sites were used for RT-PCR amplification. The resulting products were separated on 2% E-Gel agarose gels containing SYBR Safe DNA Gel Stain (Invitrogen, Waltham, MA). Bands of the expected sizes were excised, purified with the QIAquick Gel Extraction Kit (Qiagen, Germany), and subjected to Sanger sequencing on an ABI 3730 PRISM DNA Analyzer at the KI Gene – Genetic Analysis and Spatial Biology Core at Karolinska Institutet (Stockholm, Sweden).

#### *Cell fractionation*

HDFa cells were subjected to cell fractionation using two kits: the PARIS Kit (Thermo Fisher Scientific) to separate the nucleus from the cytoplasm, and the Mitochondria Isolation Kit for

Cells (Thermo Fisher Scientific) to isolate mitochondria from the cytosol, both according to the manufacturers' instructions. RNA was then extracted from each fraction.

***RNA extraction and qRT-PCR***

RNA extraction from HDFa cells was performed using TRIzol reagent (Thermo Fisher Scientific). For skin samples, the dermis and epidermis were first separated using Dispase. The dermis was sectioned with a cryotome and homogenized using a TissueLyser LT (Qiagen, Germany). Total RNA was then extracted with the RNeasy Fibrous Tissue Kit (Qiagen) according to the manufacturer's instructions. Reverse transcription was carried out using the RevertAid First Strand cDNA Synthesis Kit (Thermo Fisher Scientific). Gene expression was quantified using TaqMan or SYBR Green assays (Thermo Fisher Scientific)

***Plasmid construction***

CircMT-RNR2 overexpression plasmid was constructed using the method as previously described (*2*). Briefly, full length of circMT-RNR2 together with its upstream and downstream flanking fragments were subcloned into the pcDNA3.1 vector. The plasmid sequence was confirmed by Sanger Sequencing.

***Seahorse assay***

The oxygen consumption rate (OCR) was measured using the XF24 extracellular flux analyzer (Seahorse Bioscience, Chicopee, MA) with the Mito Stress Test (Agilent Technologies, Santa Clara, CA). HDFa cells were seeded at 20,000 cells per well in XF24-well plates (Agilent Technologies) containing 200 μL of culture medium and incubated at 37 °C with 5% $CO_2$ for 24 hours. For OCR measurement, the culture medium was replaced with 100 μL of XF assay medium (Agilent Technologies) supplemented with 2 mM glutamine (Sigma-Aldrich), and cells were incubated for 1 hour in a $CO_2$-free incubator during probe cartridge calibration. Drugs were sequentially loaded into the cartridge as follows: oligomycin (1 μM) in port A (Sigma-Aldrich), FCCP (3 μM) in port B (Tocris), and rotenone (0.5 μM) plus antimycin A (0.5 μM) in port C (both from Merck). After calibration, the cell plate was inserted and basal OCR recorded. Data were normalized to protein content.

***Protein isolation and Western Blot***

Protein lysates from HDFa cells were extracted using radioimmunoprecipitation assay (RIPA) buffer (ThermoFisher Scientific) supplemented with protease inhibitors. Protein concentrations were determined using the BCA Protein Assay Kit (ThermoFisher Scientific). Equal amounts of protein were separated on TGX precast gels (Bio-Rad, Hercules, CA) and transferred to nitrocellulose membranes. Antibody dilution ratios are listed in **Table S8**. Protein band intensities were quantified using Image Lab software (Bio-Rad).

***Histological and immunofluorescent staining***

Mouse tissue sections were processed at the Histology Core, Karolinska Institute. After dehydration, tissues were embedded in paraffin and sectioned using a microtome. For histological analysis, slides were stained with Hematoxylin and Eosin (H&E) following the manufacturer's instructions (Histolab, Korea) and imaged in brightfield mode using a Nikon Eclipse Ni-E fluorescence microscope. Wound contraction and re-epithelialization were quantified using ImageJ.

For immunofluorescent staining, formalin-fixed, paraffin-embedded tissue sections from human *ex vivo* or mouse *in vivo* wounds were deparaffinized and subjected to heat-induced antigen retrieval in Tris-EDTA buffer (pH 9) at 98 °C for 25 minutes. Sections were then blocked with 5% bovine serum albumin (BSA; ThermoFisher Scientific) and incubated overnight at 4 °C with the primary antibody (Table S8). After washing with TBST (TBS with 0.1% Triton X-100), sections were incubated with the appropriate secondary antibody for 1 hour at room temperature. Following PBS washes, slides were mounted using ProLong™ Diamond Antifade Mountant with DAPI (ThermoFisher Scientific). Immunofluorescence images were captured at 20× magnification using the Nikon Eclipse Ni-E fluorescence microscope. Images were quantified by Image J.

***Silver staining***

Silver staining was performed on the pulldown products, which were separated by SDS-PAGE and stained using the Pierce Silver Staining Kit (Invitrogen, ThermoFisher Scientific) according to the manufacturer's protocol.

***Cellular Thermal Shift Assay (CETSA) and ImmunoBlot***

HDFa cells were transfected with 60 nM si-CircMT-RNR2 or si-Control for 24 hours. After incubation, cells were rinsed and pelleted in PBS. To ensure equal cell density across conditions, cells were counted prior to lysis. Lysis was performed using NP40 buffer supplemented with protease inhibitors (Roche, Switzerland) and SUPERase RNase inhibitor (Thermo Fisher Scientific). Lysates were centrifuged at 15,000 × g for 20 minutes at 4°C, and the resulting supernatants were aliquoted into PCR tubes. Samples were then subjected to heat treatment for 3 minutes at temperatures ranging from 37°C to 60°C using a ProFlex Gradient Thermal Cycler (ThermoFisher Scientific), followed by cooling to room temperature. Protein detection was carried out with an anti-human PRDX3 antibody (1:50; ab222807, Abcam) and analyzed on the ProteinSimple Jess/Wes capillary-based system (Bio-Techne, Minneapolis, MN) according to the manufacturer's instructions.

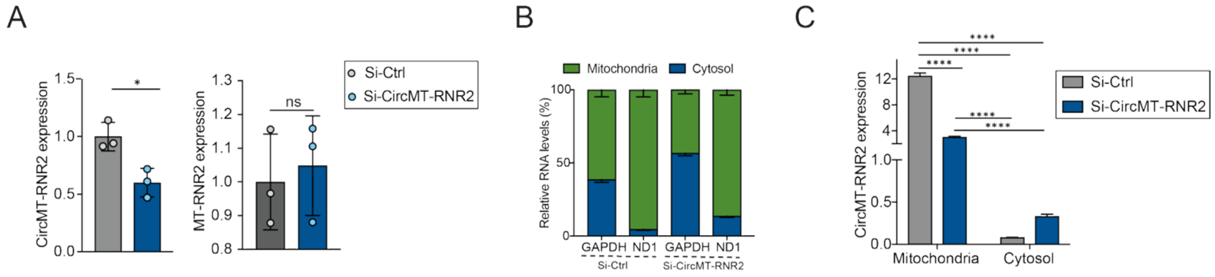


**Supplementary Figure 1: CircMT-RNR2 knockdown in human dermal fibroblasts. (A)** qRT-PCR of circMT-RNR2 and MT-RNR2 expression in HDFa cells with circMT-RNR2 knockdown (n=3). (**B–C**) qRT-PCR of ND1, GAPDH, and circMT-RNR2 in mitochondrial and cytosolic fractions of HDFa with circMT-RNR2 knockdown (n=3). ns, P≥0.05, *P<0.05, ****P<0.0001( unpaired Student t.test, or one way ANOVA and Tukey’s multiple comparison test).

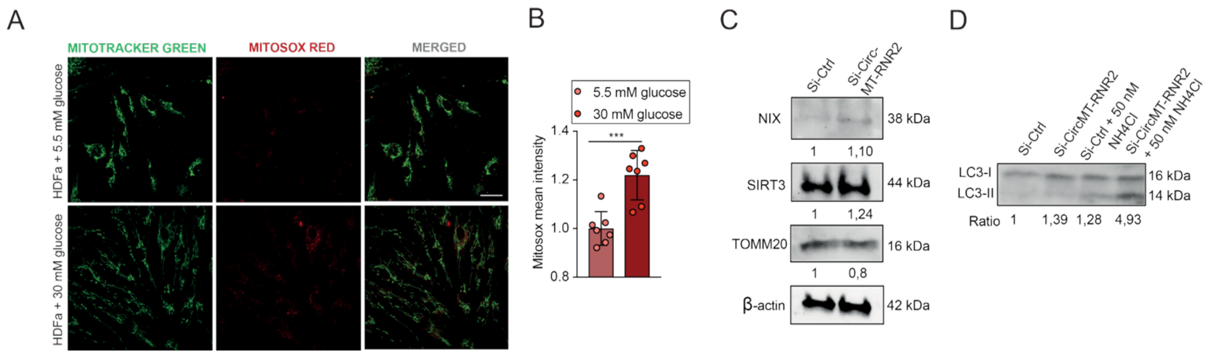


**Supplementary Figure 2. Oxidative stress under hyperglycemia and the effect of circMT-RNR2 knockdown on mitophagy. (A)** Representative confocal images of HDFa cells cultured in low (5.5 mM) or high (30 mM) glucose and stained with MitoTracker and MitoSOX (n=7); scale bar, 30 µm. **(B)** Quantification of MitoSOX Red mean fluorescence intensity from **(A)**. **(C)** Western blot of mitophagy markers (NIX, SIRT3) and mitochondrial integrity marker (TOMM20) in HDFa cells transfected with siCtrl or si-circMT-RNR2, with band quantification. β-actin served as a loading control. **(D)** Western blot of autophagy marker LC3 in circMT-RNR2 knockdown HDFa cells with or without $NH_4Cl$ (50 nM), with quantification of LC3-II/LC3-I ratio. ***P < 0.001 (unpaired Student t-test).

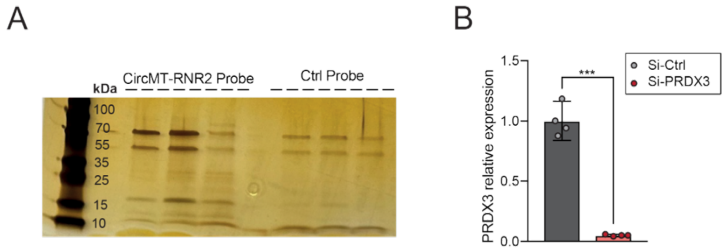


**Supplementary Figure 3. CircMT-RNR2 interacts with and stabilizes PRDX3. (A)** Silver staining of RNA pulldown products using circMT-RNR2 or control probes. **(B)** qRT-PCR analysis of PRDX3 expression in HDFa cells transfected with PRDX3 or control siRNAs (n = 4). *P < 0.001(unpaired Student t-test).

A

| Mice characteristics (Day 0) | | | | | |
|---|---|---|---|---|---|
| WT mice | Glucose levels(mmol/l) | Weight (mg) | db/db mice | Glucose levels(mmol/l) | Weight (mg) |
| 1 | 9,6 | 24,8 | 1 | 13,3 | 43,1 |
| 2 | 5,1 | 23,2 | 2 | 25,1 | 48,8 |
| 3 | 9,7 | 26,9 | 3 | 17,8 | 50,3 |
| 4 | 6,3 | 25,6 | 4 | 17,2 | 51 |
| 5 | 7,4 | 27,8 | 5 | 18,8 | 50,2 |
| 6 | 10 | 23,3 | 6 | 30 | 50,8 |
| 7 | 9,2 | 26,9 | 7 | 23,8 | 50,2 |
| 8 | 6,4 | 24,3 | 8 | 14,7 | 49,4 |
| 9 | 7,4 | 23,3 | 9 | 11,1 | 45,5 |
| 10 | 8,6 | 25,9 | 10 | 30 | 48,9 |
| 11 | 5,7 | 22,5 | 11 | 18,5 | 44,3 |
| 12 | 6,9 | 23,8 | 12 | 23,1 | 50,3 |
| 13 | 7,9 | 24,4 | 13 | 20,7 | 46 |
| 14 | 7,8 | 24,6 | 14 | 15,7 | 44,5 |
| 15 | 7,4 | 26 | 15 | 22,8 | 51,6 |

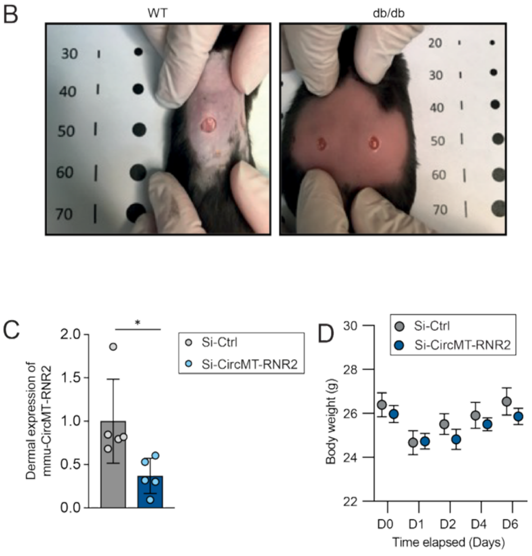


**Supplementary Figure 4. CircMT-RNR2 is required for wound closure *in vivo*. (A)** Blood glucose and body weight of wild-type (WT) and db/db mice (n = 15/group) on Day 0. **(B)** Representative wound images from WT and db/db mice. **(C)** qRT-PCR of mmu-circMT-RNR2 expression in dermal tissue from Day 6 wound biopsies treated with si-CircMT-RNR2 or control siRNA (n=5). **(D)** Body weight changes from Day 0–6 after siRNA treatment (n=5). *P < 0.05 (unpaired Student t-test).

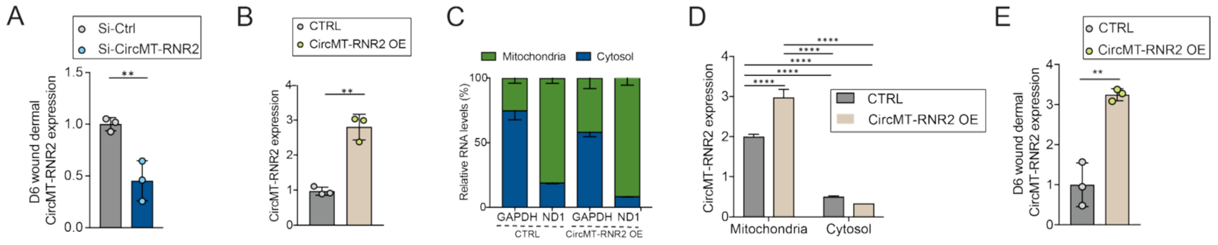


**Supplementary Figure 5: Modulation of circMT-RNR2 expression in human dermal fibroblasts and *ex vivo* wounds. (A)** qRT-PCR analysis of circMT-RNR2 in dermis from *ex vivo* wounds after 6 days of si-circMT-RNR2 treatment (n=3). **(B)** qRT-PCR analysis of circMT-RNR2 in HFDa cells with circMT-RNR2 overexpression (OE, n=3). (**C–D**) qRT-PCR of ND1, GAPDH, and circMT-RNR2 in mitochondrial and cytosolic fractions of HDFa with circMT-RNR2 OE (n=3). **(E)** qRT-PCR of circMT-RNR2 in dermis from *ex vivo* wounds after 6 days of circMT-RNR2 OE (n=3). *P < 0.05; **P < 0.01; ****P < 0.0001 (unpaired Student t-test, or one way ANOVA and multiple comparison).
